# Innate Immune Function of Neutrophil Cytoplasts Generated Post-Vital NETosis

**DOI:** 10.64898/2026.02.06.704503

**Authors:** Nithish Raj Prasad, Balaji Ganesh, Steven Dudek, Chinnaswamy Tiruppathi, Asrar B. Malik

## Abstract

The migration of polymorphonuclear neutrophils (PMN) into the site of infection such as lungs during pneumonia is a canonical feature of innate immunity. Formation of neutrophil-derived extracellular traps (NETs), web-like strands of varying lengths comprising DNA, histones, elastase, and myeloperoxidase, is an important determinant of PMN-mediated innate immunity. NETs form in microvessels, entrap bacteria and effete matter, and dampen PMN-mediated inflammatory injury at specific sites. However, studies have largely focused on NET release secondary to lytic NETosis and lysis of PMN. Far less is known about vital NETosis occurring in the absence of PMN rupture. As vital NETosis is characterized by generation of anuclear PMN termed cytoplasts (PMN_cyto_), we addressed the function of PMN_cyto_ as a critical determinant of PMN-mediated innate immunity. Studies were made in mice challenged with live *Pseudomonas aeruginosa* (PA) i.t. to induce fulminant pneumonia characterized by tissue injury in which we determined the role of generated PMN_cyto_ population. Using Tomato Red (tDTomato) transgenic mice to mark PMN, we observed PA pneumonia induced PMN transmigration leading to PMN_cyto_ generation in the airspace. In contrast, PMN_cyto_ transmigration, was minimal. PMN_cyto_ accumulating in lung tissue actively phagocytosed and killed PA. Instillation of *ex vivo* generated PMN_cyto_ also prevented PA-induced inflammatory lung injury and reduced mortality as compared to control mice. We demonstrated that the salutary effects of PMN_cyto_ required functional microchondria. Proteomic analysis revealed that PMN_cyto_ retained bactericidal and ROS generating pathways, consistent with an intact plasma membrane. Genetic deletion of peptidyl arginine deaminase 4 (PAD4), which mediates histone citrullination and promotes NETosis, facilitates PMN_cyto_ generation and thereby abrogated pneumonia-induced mortality. Thus, we have identified the crucial host defense function of PMN_cyto_ generated post-vital NETosis, suggesting that PMN_cyto_ hold promise as cell based anti-bacterial therapy in pneumonia.

## Introduction

Polymorphonuclear neutrophils (PMN), constituting 50 to 70% of all circulating leukocytes in humans, play an obligatory role in innate immunity and host defense necessary for survival (Hackert et al., 2023). PMN have been defined as the body’s first line of defense against invading pathogens because they rapidly migrate and release a repertoire of antimicrobial peptides, proteases (e.g., elastase (NE)), and enzymes (e.g., myeloperoxidase (MPO)) (Ali et al., 2025; Koenderman & Vrisekoop, 2025; Wigerblad & Kaplan, 2023; Zhang et al., 2024). PMN also executes antimicrobial defense pathways through activation of phagocytosis (Nauseef & Borregaard, 2014), degranulation of several proteases (Zhang et al., 2024), generation of reactive oxygen species (ROS), and unfettered release of anti-inflammatory cytokines (Gideon et al., 2019; Ofori et al., 2023). In addition, PMN transmigrate through endothelial adherens junctions and undergo priming at the site of infection (Mukhopadhyay et al., 2024). PMN also form extracellular traps (NETs), web-like intravascular structures comprising DNA, chromatin, and elastases in microvessels (Brinkmann et al., 2004). NETosis involves activation of peptidylarginine deiminase 4 (PAD4) required for citrullination of histones, and chromatin decondensation and nuclear content extrusion(D’Cruz et al., 2018). This process also involves release of MPO, a PMN-expressed enzyme to disassemble nucleosomes by interacting with DNA (Sollberger et al., 2018). There is another *pari passu* poorly understood, distinct from lytic NETosis, referred to as vital NETosis (Brinkmann et al., 2004; Pilsczek et al., 2010). Vital NETosis has been described by Kubes et al., in response to *Staphylococcus aureus-*induced skin infection (Pilsczek et al., 2010; Yipp et al., 2012). PMN_cyto_ generation was shown to be ROS-independent and required TLR2 signaling (Yipp et al., 2012). However, little is known about the function of PMN_cyto_ beyond their identification as anuclear cells. Thus, studies were made in the mouse model of *Pseudomonas aeruginosa* (PA) pneumonia in which infection was confined to the airspace (Liu et al., 2011). We observed generation of PMN_cyto_ post-vital NETosis largely in the airspace. Intratracheal instillation of *ex vivo* generated PMN_cyto_ killed PA and prevented pneumonia-induced lung injury, and significantly reduced mortality. PMN_cyto_ mediated host defense function was dependent on functioning mitochondria, which waned over time. However, the defective PMN_cyto_ could be restored by mitochondrial transfer. Thus, vital NETosis and the generation of PMN_cyto_ may serve as a crucial host defense mechanism for restoring homeostasis during lung infection.

## Results

### PMN_cyto_ generation post-vital NETosis

We first studied the kinetics of PMN_cyto_ generation using live confocal imaging (Zeiss LSM 710 BiG 2 GaAsP detector microscope) in transgenic Catchup mice in which PMN are genetically labelled with Tomato red gene (tDTomato) (Hasenberg et al., 2015). In addition, DNA was stained in blue by SYTO40 (Whiddon et al., 2019) to identify the loss of DNA in PMN_cyto_. In initial proof-of-concept studies, isolated mouse PMN were challenged with LPS (100µg/mL) to define features of vital NETosis in culture. We observed DNA extrusion within 10m in response to low dosage LPS challenge. The plasmalemma remained intact (**Figure 1A**) as has been described by Kubes. This finding is consistent with the definition of vital NETosis (Pilsczek et al., 2010). Many PMN in this experiment retained characteristic polymorphic nuclei and co-existed with the generated anuclear PMN_cyto_ **(Figure 1B)**. Bar graph at right shows relative number of PMN_cyto_ and normal appearing PMN FOV at 0 time (baseline) and at 30m. While PMN_cyto_ number was reduced over time relative to PMN, nevertheless, PMN_cyto_ accumulation remained high.

**Figure 1.**
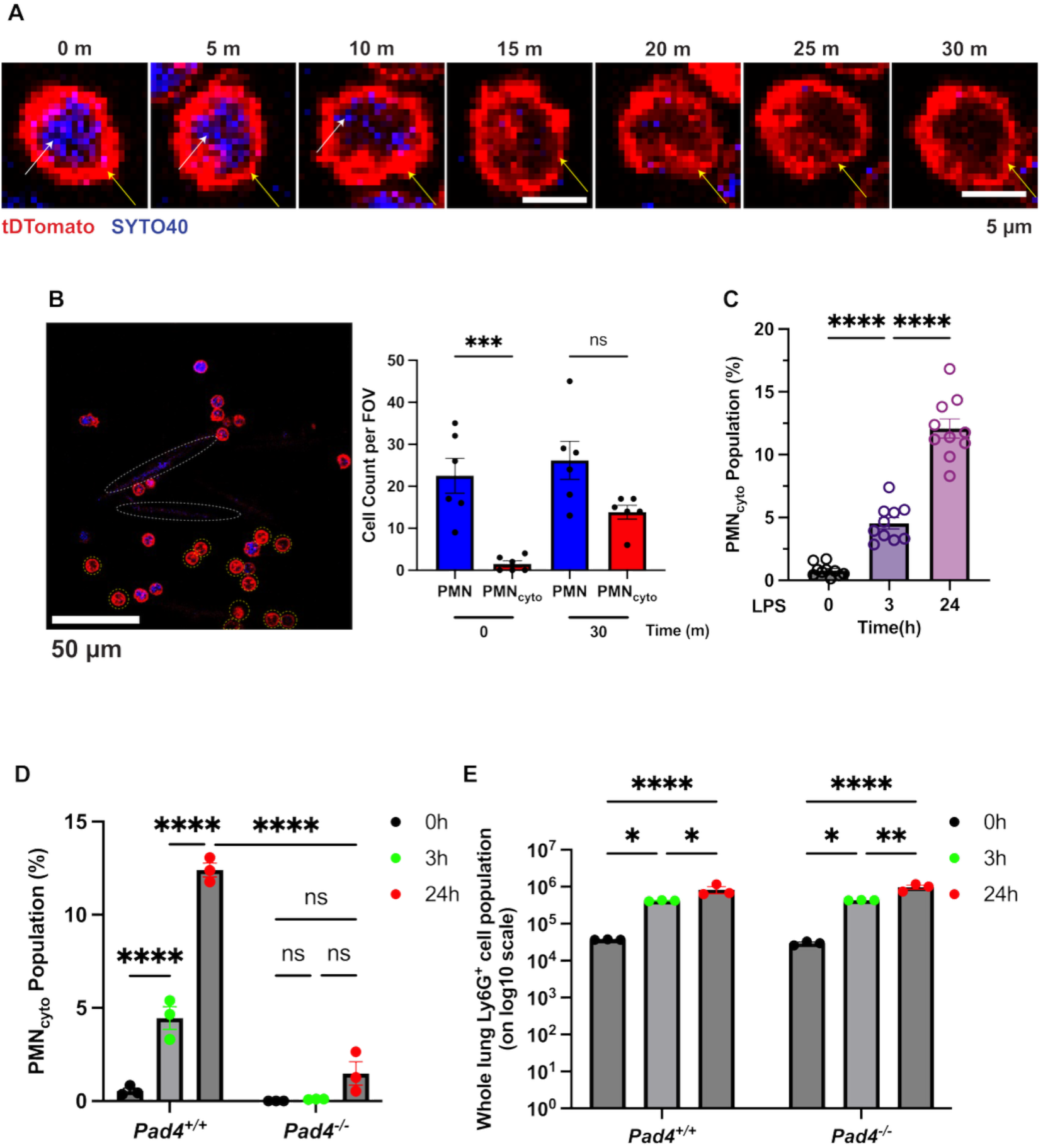
PMN_cyto_ generation post-vital NETosis requires histone citrullinating by enzyme peptidyl arginine deaminase-4 (PAD4) in mice. **(A)** PMN_cyto_ generation was detected using live confocal imaging. Images show time course of PMN_cyto_ formation determined in transgenic Catchup mice in which PMN are genetically stained. DNA staining was simultaneously assessed by SYTO40 to detect extrusion of DNA, which showed markedly reduced in blue stain (white arrow) within 10 minutes after onset of endotoxemia, and this occurred in the absence of frank rupture of PMN plasmalemma (yellow arrow) as evident by Tomato red staining. Scale bar=5µm. **(B)** Live imaging of PMN isolated from Catchup transgenic mice challenged with LPS (100 µg/ml) with DNA stained using SYTO40 and PMN in tDTomato mice. PMN_cyto_ that had not undergone lytic NETosis were identified by absence of DNA and nucleii but retained plasmalemma. Bar graph at right shows relative number PMN_cyto_ and normal PMN both before and after LPS (100µg/mL) challenge. Blue reflects DNA staining with SYTO40 released post NETosis. Shown are mean ± SEM, n=5, ns= not significant, ***p<0.0001. **(C)** Time course of PMN_cyto_ generation in mice as determined by FACS analysis of lung single cell suspensions obtained post LPS. Cells stained with DNA marker SYTO40 and PMN marker AF647-Ly6G mAb were used to determine relative number PMN_cyto_ generated. PMN_cyto_ appeared as early as 3h after onset of endotoxemia and continued to expand over next 24h ascribable to vital NETosis. N=10; ****p<0.001, shown are mean ± SEM. **(D)** The enzyme peptidyl arginine deaminase 4 (PAD4), which citrullinates histone, is required for PMN NETosis. Studies were made in *pad4^-/-^* and WT mice i.p. injected with LPS, cell suspensions were used for FACS analysis to determine the relative number of PMN and PMN_cyto_ in lungs. Results show marked PMN_cyto_ generation in WT mice and severely defective response in *Pad4^-/-^* mice. N=3, **p<0.01, mean±SEM. **(E)** Deletion of PAD4 *per se* had no independent effect on accumulation of Ly6G+ PMN indicating that inhibited PMN_cyto_ generation in *Pad4^-/-^* mice (shown in Figure 1D) cannot be ascribed to differences in relative number of lung PMN sequestration between WT and *Pad4^-/-^*mice.

In addition, in mouse studies we observed that i.p LPS (10mg/kg) increased PMN_cyto_ generation over 24h **(Figure 1C)** indicating generation of PMN_cyto_ post endotoxemia. To further address the PMN_cyto_ generation *in vivo,* we determined the role of peptidyl arginine deaminase 4 (*pad4*), the enzyme that citrullinates histones and is required for NETosis (Li et al., 2010). Thus, studies were made in *Pad4^-/-^* mice to determine whether preventing NET formation interfered with PMN_cyto_ generation. Results showed marked PMN_cyto_ generation in WT mice and severely defective response in *Pad4^-/-^* mice (**Figure 1D)**. However, deletion of PAD4 had no effect on accumulation of Ly6G^+^ cells **(Figure 1E)**, indicating that inhibition of PMN_cyto_ generation was independent of differences in PMN sequestration in WT and *Pad4^-/-^* mice.

### *Pseudomonas aeruginosa* (PA) pneumonia induces PMN_cyto_ airspace accumulation in mice

We next used an established PA pneumonia model in mice (Damron et al., 2016; Liu et al., 2011) to study PMN_cyto_ generation and function *in vivo*. PA was instilled into the airway with either 10^6^ or 10^7^ CFU/mouse, and NET formation was assessed using bronchoalveolar lavage fluid (BALF) by measuring PMN elastase as a real-time NETosis marker (Xiao et al., 2021). We observed PA dose-dependent NETosis (**Figure 2A**). We also observed time-dependent generation of PMN_cyto_ in lungs at 3h and up to 6h post-PA, mirroring the time course of NETosis (**Figure 2B**).

**Figure 2.**
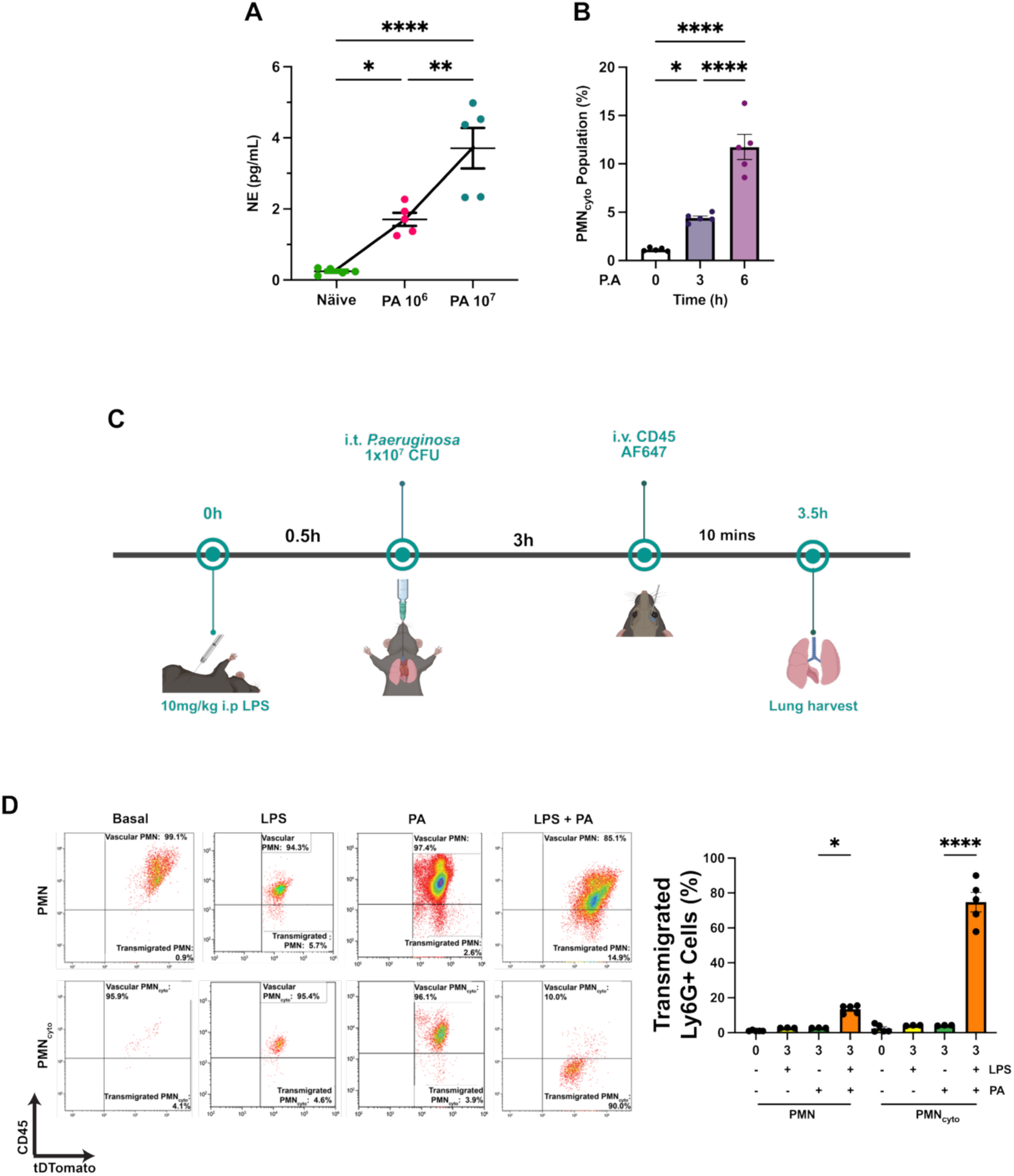
Accumulation of PMN_cyto_ in the airspace in response to *Pseudomonas aeruginosa* (PA) induced pneumonia. (**A**) WT mice were i.t. instilled with PA (either 10^6^ or 10^7^ CFU/mouse) for 24h, and NET generation was determined using bronchoalveolar fluid (BALF) by ELISA of PMN elastase (NE). NE is a surrogate marker of NETosis (Xiao et al., 2021). n=5, *p<0.05, **p<0.01, ****p<0.0001, shown are mean ± SEM. (**B**) Time-dependent increase in PMN_cyto_ number in lungs after PA infection, N=5, *p<0.05, ****p<0.0001, mean ± SEM. (**C**) Experimental scheme for the determination of PMN_cyto_ accumulation in lung airspace. Catchup mice were injected with i.p. LPS (10mg/kg) at 0 time followed by i.t. PA (1×10^6^ CFU/mouse) infection at 30m to assess PMN_cyto_ accumulation in the lung airspace. At 3h post-infection, anti-CD45 mAb was injected i.v. prior to harvesting lungs. Lung single cell suspensions were prepared, stained with SYTO40, and cells were analyzed by FACS. (**D**) Accumulation of PMN and PMN_cyto_ into lung airspace was determined as described by us (Mukhopadhyay et al., 2024). Results showed that PMN_cyto_ accumulating in airspace far exceeded PMN entry (shown on left and right). N=5, *p<0.05, ****p<0.0001, mean ± SEM.

Next to study whether PMN_cyto_ were generated locally in the airspace or accumulate in the airspace following transmigration. Here we used a model in which LPS was first i.p. injected at 0 time to generate PMN_cyto_ in the circulation, and then PA was instilled i.t. at 30m to determine PMN_cyto_ transmigration driven by airspace delivered PA (**Figure 2C**). To differentiate between transmigrated vs. intravascular PMN_cyto_, we used our method (Mukhopadhyay et al., 2024) employing Catchup mice in which PMN are genetically labeled by tDTomato, and then anti-CD45 mAb (AF67) is injected i.v. 10m before euthanasia to label PMN and any PMN_cyto_ generated in the circulation. Lungs were extracted to differentiate between transmigrated and intravascular PMN_cyto_ using FACS analysis (Mukhopadhyay et al., 2024). We observed negligible number (<1%) of PMN and PMN_cyto_ in the basal condition in the airspace (**Figure 2D**). LPS alone (i.p. 10mg/kg) also did not induce PMN_cyto_ transmigration (**Figure 2D**). Following i.t. PA (1×10^6^ CFU/mouse) for 3h, however, we observed <5% of PMN and PMN_cyto_ had transmigrated (**Figure 2D**). In contrast, LPS priming of circulating PMN markedly increased airspace PMN_cyto_ accumulation from <5% to ∼70% in response to PA pneumonia (**Figure 2D**). The FACS results showing marked enhancement of airspace PMN_cyto_ accumulation are summarized in **Figure 2D**.

### Host defense function of PMN_cyto_

We next addressed the functional significance of airspace sequestered PMN_cyto_ in the response to pneumonia. Here we adoptively transferred either PMN or PMN_cyto_ obtained from Catchup mice (1×10^4^ cells/mouse) into recipient WT (C57BL/6J (same background as Catchup mice), 30m after the mice were i.t. instilled with GFP-labelled-PA (1×10^6^ CFU/mouse) to determine whether accumulation of PMN_cyto_ interfered with *in vivo* phagocytic activity (**Figure 3A**). We observed that PMN_cyto_ phagocytosed PA similar to PMN (**Figure 3B**), and there was no difference in killing of PA by PMN_cyto_ as compared to PMN (**Figure 3C**). PMN_cyto_ also prevented lung vascular injury (a hallmark of pneumonia induced lung injury) (Bernard et al., 1994) as compared to naïve PMN **(Figure 3D)**. To address whether PMN_cyto_ improved survival, we adoptively transferred either PMN or PMN_cyto_ (1×10^4^ cells/mouse) in mice challenged with lethal dose of PA (1×10^7^ CFU/mouse) (**Figure 3E**). In these studies, PMN_cyto_ significantly improved survival by 3.5-fold compared to mice receiving PBS, and 2-fold improvement in mice receiving primed PMN (**Figure 3E**).

**Figure 3.**
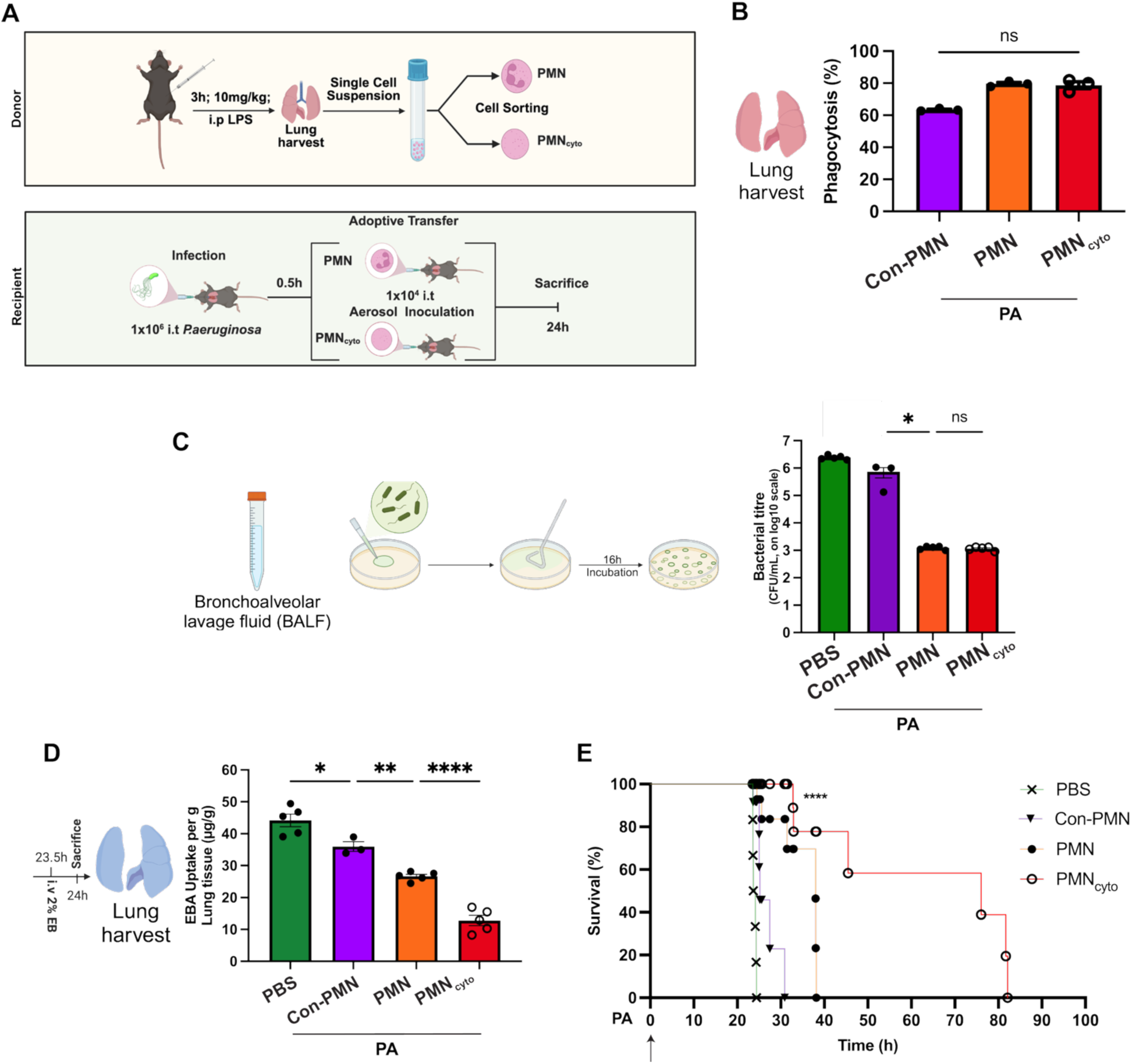
PMN_cyto_ phagocytosis of *Pseudomonas aeruginosa* (PA) and mitigation of pneumonia and inflammatory lung injury. (**A**) Schematic showing preparation of PMN and PMN_cyto_ from lungs of LPS challenged Catchup mice, and adoptive transfer of these cells into lungs of PA infected recipient WT mice. (**B**) Lungs harvested 24h after PA-infected recipient mice were used to determine phagocytosis of PA. Quantification shows PA phagocytosis induced by adoptively transferred PMN and PMN_cyto_. PMN_cyto_ demonstrated phagocytic activity equal to PMN. N=3; ns, not significant, shown are mean ± SEM. (**C**) *In vivo* bactericidal function was measured in PA infected mice using BALF collected from control mice (i), Con-PMN adoptively transferred mice (ii), PMN adoptively transferred mice (iii), and PMN_cyto_ adoptively transferred mice (iv). N=3-5, ns= not significant, *p<0.05, shown are mean values ± SEM. (**D**) Lung vascular permeability (EBA uptake in lungs; a measure of injury) was determined in each group (i-iv). Lung vascular permeability was markedly reduced in the PMN_cyto_ adoptively transferred mice compared with other groups. n=3-5, *p<0.05, **p<0.01, ****p<0.0001, mean ± SEM. (**E**) Survival curves for the 4 groups (i-iv) (1×10^4^ cells/mouse, i.t) adoptively transferred into PA (1×10^7^ CFU, i.t.) infected mice. Survival curve was compared using the Log-rank test for survival (n = 6). ****p<0.0001.

### Residual mitochondria in PMN_cyto_ limit bactericidal function

We next determined whether ATP generated by mitochondria, known to limit PMN host defense function (Weinberg & Chandel, 2025; Zmijewski et al., 2008), is a factor in attenuating the bactericidal activity of PMN_cyto_. Thus, we first analyzed any functional differences of PMN and PMN_cyto_ formed during the early phase of endotoxemia (3h) to later phase (24h) using an *in vitro* bactericidal assay. PMN and PMN_cyto_ obtained at 3h both showed similar bacterial killing, whereas PMN and PMN_cyto_ obtained at 24h showed significantly reduced bactericidal activity as compared to the 3h PMN and PMN_cyto_ (**Figure 4A**). Interestingly, the decrement in PMN_cyto_ mediated bacterial killing at 24h was significantly greater than PMN mediated bacterial killing (**Figure 4A**). Thus, we next addressed whether the attenuated response in PMN_cyto_ was the result of mitochondrial dysfunction and waning ATP generation over time. In control experiment, we demonstrated no difference in mitochondrial number of PMN vs. PMN_cyto_ groups after LPS (**Figure 4B**). Next, to address mitochondrial function, we examined mitochondrial membrane potential measurements using Tetramethyl rhodamine methyl ester (TMRM) (Ward et al., 2007). Results showed a time-dependent decrease in PMN_cyto_ mitochondrial membrane potential over the 24h period, unlike in PMN (**Figure 4C**). Next, Seahorse assay measurements also showed a temporal decrease in ATP production in PMN_cyto_, which was greater than seen in PMN (**Figure 4D**). These findings show a time-dependent decrease in mitochondrial function in PMN_cyto_.

**Figure 4.**
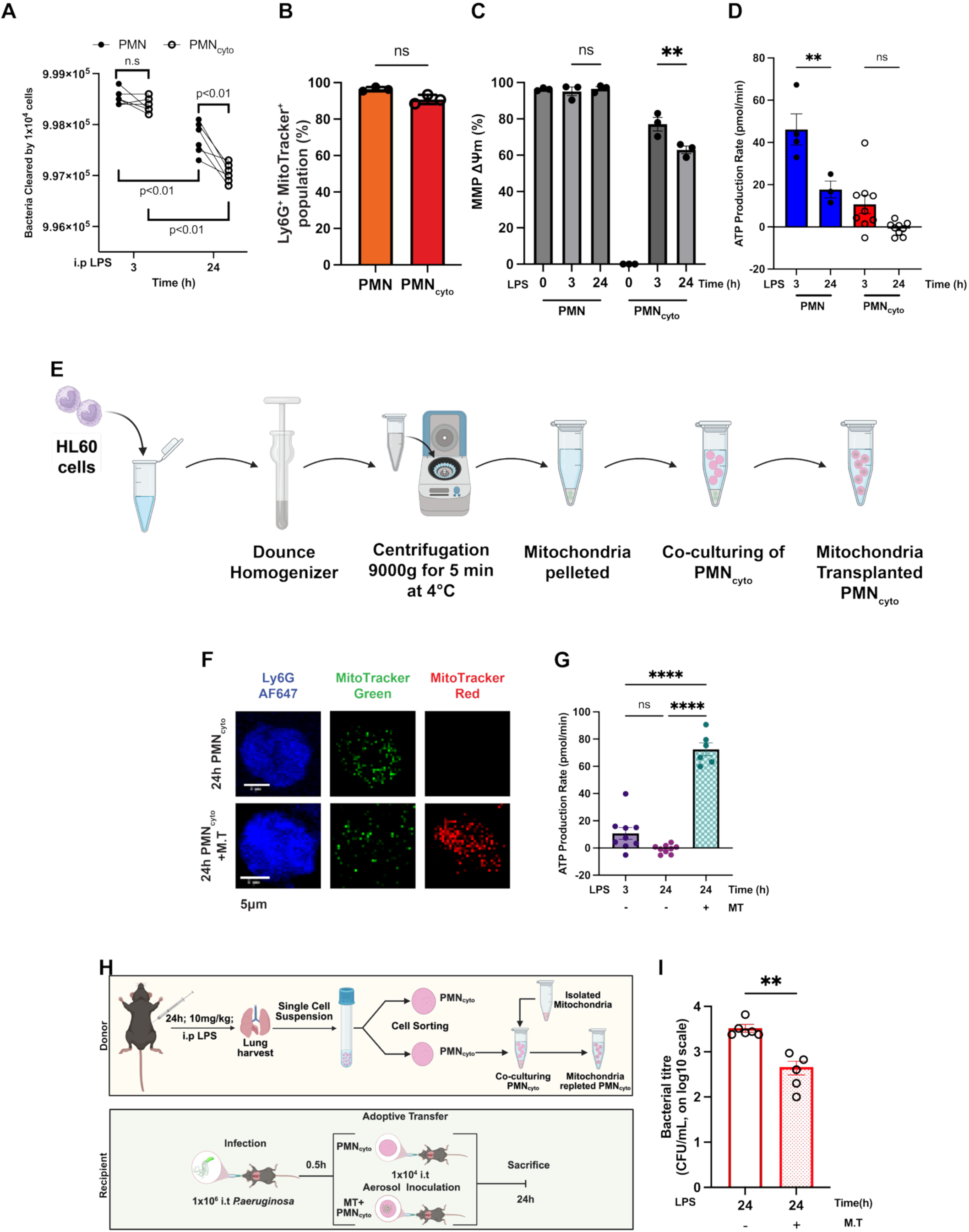
PMN_cyto_ mitochondria dysfunction limits host defense and restoring protection by mitochondria repletion. (**A**) PMN_cyto_ generated *in vivo* shows bactericidal activity similar to PMN in response to live PA. WT mice were i.p. injected with LPS for 3h and 24h, and thereafter, PMN and PMN_cyto_ were prepared and used for bacterial killing assay. N=6, ns= not significant, shown are mean ± SEM. (**B**) Catchup mice challenged with LPS for 3h and then lung PMN and PMN_cyto_ (with tD Tomato Ly6G-expressing, red) were prepared, stained with SYTO40 (DNA, blue) and MitoTracker (green) were used for FACS analysis. n=3; ns, not significant. Shown are mean ± SEM. (**C**) PMN_cyto_ mitochondrial membrane potential decreses in a time-dependent manner after LPS challenge. The mitochondrial membrane potential (MMP ΔΨm) was measured using lung PMN and PMN_cyto_ obtained from mice i.p. LPS injected for 0, 3, and 24h. n=3; ns= not significant, **p<0.01. Shown are mean ± SEM. (**D**) Mitochondria ATP generation was determined in lung PMN and PMN_cyto_ obtained from WT mice challenged with LPS for 3h and 24h using Seahorse analyzer. n=3-9; ns= not significant; **p<0.01, shown are mean ± SEM. (**E**) Schematic of procedure for mitochondria isolation (from HL60 cells) and transfer into PMN_cyto_ obtained from WT mice i.p. injected with LPS for 24h. (**F**) Confocal images demonstrate effective transfer of mitochondria into PMN_cyto_. In this experiment, PMN_cyto_ were sorted and stained with Ly6G mAb-AF647 (blue), WT mitochondria were stained with MitoTracker Green (green), and isolated mitochondria stained with MitoTracker Red (red). Scale bar= 5µm. (**G**) ATP generation as measured by Seahorse analyzer in PMN_cyto_ obtained from mice exposed to LPS for 24h (time of maximum decrease in PMN_cyto_ mitochondrial ATP production shown in Figure 4D). Results show that mitochondria transfer markedly increased ATP production in PMN_cyto_. N=6-8, ****p<0.0001, shown as mean ± SEM. (**H**) Schematic of experiment showing donor WT mice exposed to LPS for 24h from which PMN_cyto_ were obtained. Mitochondria transferred into PMN_cyto_, were adoptively transferred into PA-infected mice. (**I**) Lung bacterial killing efficiency of mitochondria repleted PMN_cyto_ demostrasted 10-fold increase as compared to 24h PMN_cyto_ in absence of mitochondrial transfer. N = 5; **p<0.01, shown are mean ± SEM.

Thus, we next addressed whether restoring time-dependent PMN_cyto_ mitochondrial dysfunction through mitochondrial transfer (Spees et al., 2006) normalizes PMN_cyto_ bactericidal activity. Mitochondria were isolated from HL60 cells and co-cultured with PMN_cyto_ from mice challenged with LPS for 24h (Wigler & Weinstein, 1975) (**Figure 4E**). To differentiate between native and transferred mitochondria, PMN_cyto_ mitochondria were labeled with MitoTracker green, and HL60-derived mitochondria were labeled with MitoTracker red. We observed efficient transfer of MitoTracker Red-labeled mitochondria into mitochondria-depleted PMN_cyto_ (**Figure 4F**). ATP production showed a significant increase in mitochondria transferred (MT) PMN_cyto_ (**Figure 4G)**. Next to study whether host defense function of mitochondria-repleted PMN_cyto_ was restored, we adoptively transferred 1×10^4^ mitochondria-transferred PMN_cyto_ into WT mice, and compared PA response with mitochondria-depleted PMN_cyto_ (**Figure 4H**). Bactericidal activity was studied after i.t. instillation with PA (1 × 10^6^ CFU/mouse) followed by i.t. mitochondria-transferred PMN_cyto_ in mice. Importantly, mitochondria-transferred PMN_cyto_ mice normalized bacterial clearance as compared with mitochondria-depleted PMN_cyto_ control mice (**Figure 4I)**.

### Proteomic profiling of PMN_cyto_ shows persistence of host defense machinery

Proteomic analysis was carried out to determine whether PMN_cyto_ generated following loss of nuclei DNA retained the host defense machinery of PMN. Lung single cell suspensions were obtained from Catchup mice challenged with LPS for 3h, followed by cell sorting, by staining with nuclear marker SYTO40. Using mass spectrometry, we found expression of 4571 proteins in PMN and 3392 proteins in PMN_cyto_ (**Figure 5A**). We also observed exclusive expression of 1357 proteins in PMN and 178 proteins in PMN_cyto_ (**Figure 5A**). Key proteins associated with NETosis, such as MPO (Myeloperoxidase) and ELANE (Neutrophil Elastase), were expressed in both groups. Surface markers of PMN, such as CD177, CD11b, and CD18, were, however, expressed at low levels in PMN_cyto_ as compared to PMN (**Figure 5B, C**). Functional GO pathway analysis showed enhanced expression of chemoattractant and inflammatory regulatory pathway proteins in PMN_cyto_ as compared with PMN (**Figure 5D**). Furthermore, expression of mitochondrial ETC and glycolytic pathway-specific proteins were not different between PMN and PMN_cyto_ (**Figure 5D).**

**Figure 5.**
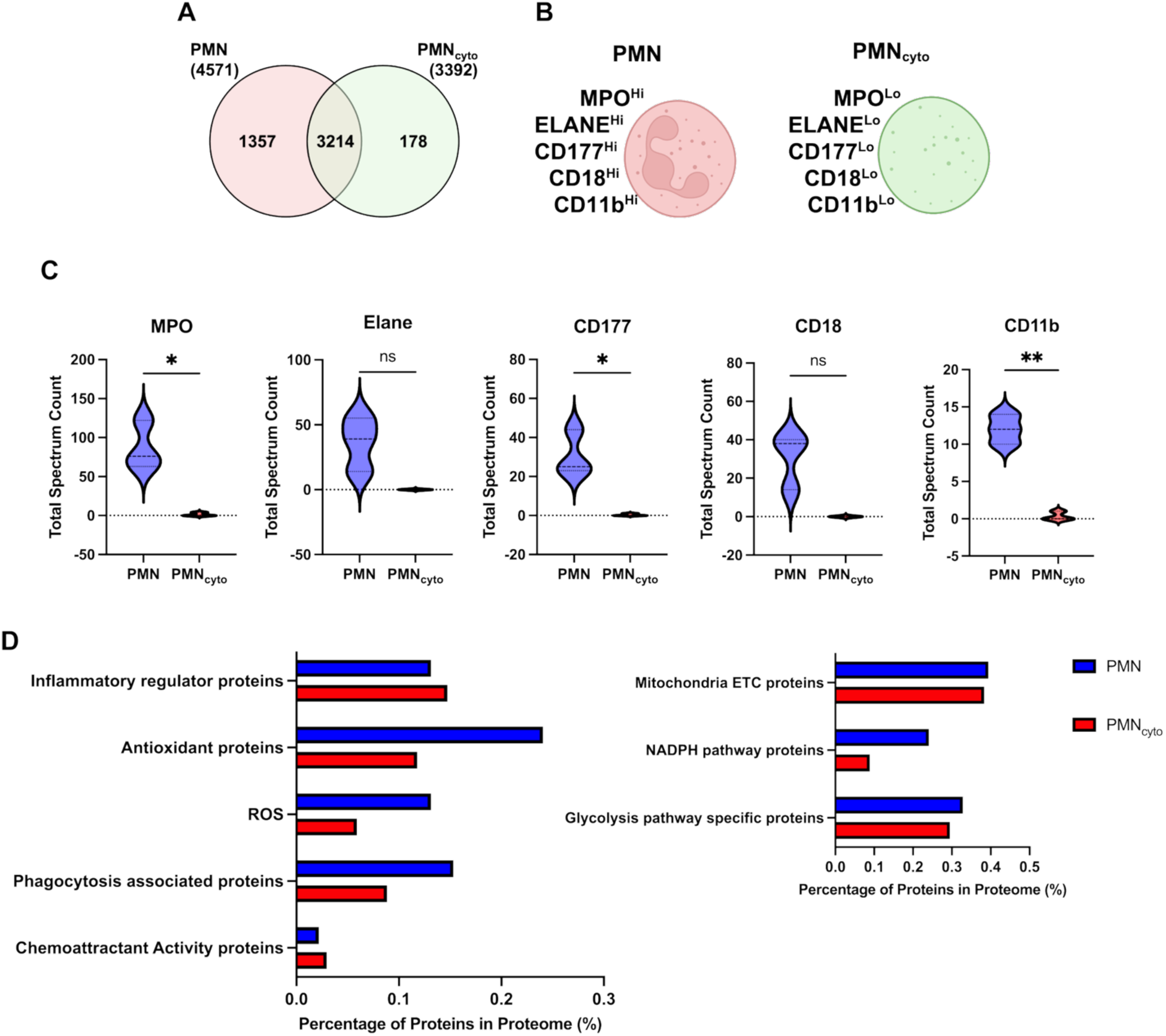
Mass spectrometric identification of protein expression pattern in PMN and PMN_cyto_: Catchup mice were challenged with LPS for 3h, then PMN and PMN_cyto_ were separated by cell sorting after staining with nuclear marker SYTO40, and the cells were used for mass spectrometry. (**A**) Venn diagram shows the expression of total number of proteins, unique and overlapping proteins, in PMN and PMN_cyto_. (**B-C**) Proteins that distinguish PMN and PMN_cyto_ (**B**) and their expression levels (**C**). N = 3; ns= not significant; *p<0.05, **p<0.01, shown are mean ± SEM. (**D**) GO pathway protein enrichment in PMN and PMN_cyto_. Results shown are mean values of 3 independent experiments.

## Discussion

NET formation in microvessels secondary to PMN sequestration and activation entraps bacteria and promotes resolution of inflammatory tissue injury (Brinkmann et al., 2004). This host defense function is among the cardinal PMN innate immune mechanisms protecting the host (Beiter et al., 2006). In the present study, we identified another previously unrecognized protective pathway engaged via the generation of an anuclear population of PMN_cyto_ secondary to the phenomenon referred to as vital NETosis (Pilsczek et al., 2010). These studies identified PMN_cyto_ formation in mice challenged with *S.aureus* (Pilsczek et al., 2010), consistent with our observation following *Pseudomonas aeruginosa*-induced pneumonia. Despite evidence of their formation *in vivo,* little is known about the function of these cells, such as whether they retain PMN-like characteristics and whether they have a survival advantage for the host. Intriguingly, our results showed that PMN_cyto_ demonstrated a potent bactericidal activity in the mouse pneumonia model (Liu et al., 2011). We showed robust generation of PMN_cyto_ *ex vivo* using naïve human (**Supp. Figure 1**) or mouse PMN (**Figure 1A,B)** used in these studies. We also showed that PMN_cyto_ survived up to 2 weeks, enabling assessment of their function (**Supp. Figure 2**).

Results showed that instillation of PMN_cyto_ in mice with established *PA* pneumonia prevented lung injury and reduced mortality, and the protection was secondary to enhanced *PA* killing by PMN_cyto_. Naïve PMN, similarly instilled in the same number, also showed protection, although it was less than PMN_cyto_. Thus, the present studies demonstrate the protective action of PMN_cyto_ against *PA* pneumonia because they retained the bactericidal function of PMN.

An important observation was that PMN_cyto_ bactericidal activity gradually waned over 24h. We attributed this response to a time-dependent loss of mitochondrial function and reduced ATP generation in PMN_cyto_, and the known requirement for ATP generation for PMN bactericidal activity (Bao et al., 2014). Thus, we addressed whether bacterial killing was secondary to decreased ATP production. To test the role of reduced ATP production in PMN_cyto_, we repleted mitochondria in PMN_cyto_ using the standard incubation protocol (Huang et al., 2016; McCully et al., 2009). We observed that restoration of mitochondria in PMN_cyto_ revived their bactericidal activity, showing the requirement of mitochondria in sustaining their host-defense function.

To query whether NETosis in PMN *per se* was responsible for the generation of the protective population of PMN_cyto_, studies were made in mice genetically deleted of peptidyl arginine deaminase 4 (*Pad4^-/-^*) (D’Cruz et al., 2018; Li et al., 2010). PAD4 functions by citrullation of DNA and histone, and is required for NETosis (Franck et al., 2018; Li et al., 2010). We observed that the deletion of Pad4 in mice showed severely defective NETosis and had virtually no generation of PMN_cyto_. Moreover, both WT and *Pad4*^-/-^ mice upon LPS challenge recruited relatively equal numbers of Ly6G^+^ cells into the lung. This finding raises the interesting question whether PMN_cyto_ generation secondary to vital NETosis is more advantageous to the host than lytic NETosis generating NETs. Although we did not study lytic NETosis, it is possible that in the inflammatory milieu each mechanism operating in a context-specific manner has adaptive functions through either NET formation for bacterial entrapment or generation of PMN_cyto_ for bacterial killing.

We studied the function of PMN_cyto_ in pneumonic lungs to address the question of whether PMN_cyto_ are generated in the air space or in blood vessels post-vital NETosis. Thus, studies were made in which we determined the transmigration of PMN and PMN_cyto_ from vessels to lung parenchyma using an adaptation of our method to differentiate between cells in vascular and extra-vascular compartments (Mukhopadhyay et al., 2024). We observed that the majority of PMN_cyto_ present in the airspace after induction of pneumonia were derived from *de novo* PMN_cyto_ generation in the airspace as opposed to transmigration of PMN_cyto_ from vessels. Thus, the results suggest active airspace PMN_cyto_ generation occurs secondary to PMN transmigration in response to PA pnueumonia. Priming of circulating PMN with endotoxin was required for effective PMN transmigration into tissue prior to transitioning into PMN_cyto_, and subsequent PMN_cyto_-mediated protection.

To address the capability of PMN_cyto_ to mount an effective host defense response, we carried out a proteomic analysis of PMN_cyto_ primed with LPS. We observed that the host defense and bactericidal functions, i.e., ROS-associated proteins, inflammatory regulatory proteins, mitochondrial ETC proteins, chemoattractant proteins (and others as shown in **Figure 5D**) are present in PMN_cyto_. Thus, PMN_cyto_ retained the essential host-defense machinery, explaining the salutary effects of PMN_cyto_ treatment in *PA*-induced pneumonia in mice. It remains unclear whether the protective effects of PMN_cyto_ can be translated to human patients. Additional unaddressed questions are whether PMN_cyto_ efficacy is demonstrable in other models of bacterial lung injury, whether PMN_cyto_ are useful in drug-resistant bacterial infections, and the time window for treatment post-infection. Our results inform the last question since fresh PMN_cyto_ showed protection, which was the function of having a normal load of mitochondria as described above.

An important question is whether resident or *de novo* airspace migrating macrophages promote clearance of PMN_cyto_ through efferocytosis (Aggarwal et al., 2014; Nepal et al., 2019; Proto et al., 2018). However, our results showed that macrophages did not engulf PMN_cyto,_ and the PMN_cyto_ remained profoundly bactericidal (**Supp. Figure 3**). We also observed PMN_cyto_ in lungs persisted up to 2 weeks (**Supp. Figure 2**) after LPS challenge, further suggesting that macrophages do not play a role in clearing and limiting the function of PMN_cyto_.

In conclusion, our findings support the critical role of PMN_cyto_ in anti-bacterial host defense. We showed that PMN_cyto_ generation after vital NETosis served to dampen the severity of inflammatory tissue injury and that PMN_cyto_ could be normalized by mitochondrial transfer to prolong PMN_cyto_ bactericidal activity. Moreover, in the absence of PMN_cyto_ generation following genetic deletion of PAD4 and prevention of NETosis we observed enhanced mortality induced by *PA.* As instillation of naïve PMN in the PA pneumonia model did not reduce mortality as compared to PMN_cyto_ the results support the cell therapy potential of PMN_cyto_. Thus, generation of PMN_cyto_ serves as an intrinsic host defense mechanism to limit inflammatory injury. This may be a telologically relevant bactericidal mechanism to dampen overwhelming PMN assault and thus limit tissue injury.

## Materials and Methods

### Experimental Models

#### Animals

Mice were housed and bred under specific pathogen-free conditions at the University of Illinois, Chicago (UIC) Animal Care Facility. All experiments were performed under the ethical principles and guidelines approved by the UIC Institutional Animal Resources Center animal usage committee. Male and female mice aged between 8 and 16 weeks were used for experiments. C57BL/6 was purchased from the Jackson Laboratory, and Catchup mice were obtained from the University of Duisburg–Essen (Hasenberg et al., 2015; Mukhopadhyay et al., 2024).

#### Inflammatory lung injury model in mice

Mice were intraperitoneally (i.p.) injected with LPS (Lipopolysaccharide from *Escherichia coli* O111: B4, L2630, Lot 0000211692; Sigma-Aldrich) dissolved in PBS. *Pseudomonas aeruginosa* (PA) expressing GFP (GFP-PA01)(Joshi et al., 2020), 1 × 10^6^ /10^7^ CFU in 40µl of PBS, was intratracheally (i.t.) sprayed using micro aerosol spray (Liquid PenWu Device, BJ-PW-M; Biojane).

#### *In vitro* imaging analysis of PMN_cyto_ formation

Live imaging of *in vitro* studies was accomplished using the Zeiss LSM 710 BiG, a confocal microscope with an incubator driven by the ZEN software. ImageJ was used to quantify the cytoplast formation rate. The stained cells (DNA-SYTO40, PMN-Ly6G) were collected in a fibronectin-coated 35mm petri dish with a 14mm microwell with a No.1.5 coverglass. The cells were left undisturbed for adhesion to the fibronectin-coated (Bohnsack et al., 1990; Kuntz & Saltzman, 1997) coverglass for 20 minutes at 37°C and 5% CO_2_. After adhesion, the cells are transferred to the microscope equipped with the incubator before being challenged with 100µg/mL of LPS. It is then continuously imaged for 30 minutes, capturing a frame every 1 minute at 40x magnification. Acquired tiles were stitched using ZEN software and analyzed using ImageJ.

### Lung single cell suspension for flow cytometry

C57BL/6 mice lungs were harvested, and a single cell suspension was prepared by digestion of minced lung lobes with collagenase A (1 mg/ml; Millipore-Sigma # SCR103) at a 37°C shaking water bath for 45 min. Digested lung samples were passed through a 16G needle and filtered through a 40µm cell strainer (Thermo Fisher, #08-771-1). Red blood cells were removed by using Ter119-microbeads (Miltenyi Biotec, # 130-049-901). The cells were stained with Ly6G-AF647 (mAb) and SYTO40 ex vivo for 30 minutes.

### Flow cytometric (FACS) analysis and cell sorting

PMN, labeled with Ly6G or tdTomato, and its nucleus stained with SYTO40. CytoFLEX-S (Beckman) flow cytometer, equipped with 405, 488, 561, and 635 nm lasers, was used for data acquisition of all the flow cytometry data. Data analyses were performed using Kaluza Analysis 2.2 (Beckman) software.

PMN and PMN_cyto_ were sorted in a sterile environment by MoFlo Astrios and MoFlo Astrios EQ (Beckman Coulter) cell sorter equipped with 405, 488, 561, and 635 nm lasers and collected in RPMI-1640 medium (Thermo Fisher #72400047). Based on granularity and relative size, gates were initially established on forward and side scatter. Aggregates were gated out accordingly. Selective gates were drawn to sort the PMN and PMN_cyto_. Software SUMMIT 6.1 (Beckman Coulter) was used for sorting and analysis.

### LC/MS proteomic analysis

Catchup mice were challenged with LPS for 3h and stained for their nucleus using SYTO 40. Single cell suspension is prepared, and using FACS, PMN and PMN_cyto_ were sorted. The 2 cell type samples had their proteins extracted using RIPA buffer (Sigma R0278) plus solutions of 20x Protease Inhibitor cocktail (Sigma P2714), 100x Phosphatase Inhibitor cocktail 3 (Sigma P0044), and 100x Phosphatase Inhibitor cocktail 2 (Sigma P5726) for the final volume of 2mL. Under ice, we added 400uL of this RIPA solution with the inhibitors to the cell pellets. We used sonication-trap (QSonica Sonicator) with 5 seconds of sonication, 5 seconds of standby, 40% amplitude, for 1 minute. Afterwards, we centrifuged at 12,000 rpm for 30 minutes and obtained approximately 350μL of supernatant subjected to protein quantification and digestion with the FASP protocol. We used the BCA (Pierce BCA Protein Assay Kit) and the A280 assay to quantify the dosage of proteins present in the sample. 25μg of proteins were carried out with Filter Aided Sample Preparation (FASP) protocol, using 10K NMWL centrifugal filter units. Dithiothreitol, which was added at a final concentration of 20 mM, and reduced for 30 minutes. Proteins were then alkylated with IAA at a final concentration of 50mM IAA for 20 min in the dark. The alkylated proteins were washed with 200 µL 8M Urea in 0.1M Ammonium Bicarbonate (ABC) 3 times and equilibrated with 0.1M ABC 3 times prior to tryptic digestion. Proteins were digested using Trypsin at an enzyme to protein ratio of 1:50 in 50μL 0.1M ABC buffer at 37°C overnight. The digested peptides were collected by centrifugation at 14,000g for 20 min, followed by two elutions with 50μL of 50 mM ABC and one elution with 50μL of 0.5M NaCl. The eluted peptides were desalted using Oasis PRiME HLB (30mg) and dried. And then resuspended for injection into the Q Exactive HF. 1µg of samples were analyzed using Q Exactive HF mass spectrometer coupled with an UltiMate 3000 RSLC nanosystem with a Nanospray Frex Ion Source (Thermo Fisher Scientific). Digested peptides were loaded into a Waters nanoEase M/Z C18 (100Å, 5um, 180µm x 20m) trap column and then a 75 μm x 150mm Waters BEH C18(130A, 1.7um, 75µm x 15cm) and separated at a flow rate of 300nL/min. Solvent A was 0.1% FA in water, and solvent B was 0.1% FA, 80% ACN in water. The solvent gradient of LC was 5% B in 0-3 min, 8% B in 3.2 min, 8-35% B in 85 min, 35-95% B in 90 min, wash 95% 94.8 min, followed by 5% B equilibration until 105 min. Full MS scans were acquired in the Q-Exactive mass spectrometer over 350-1400 m/z range with a resolution of 120,000 (at 200 m/z) from 5 min to 95 min. The AGC target value was 3.00E+06 for the full scan with a 445.12003 lock mass. The 15 most intense peaks with charge states 2, 3, 4, 5 were fragmented in the HCD collision cell with a normalized collision energy of 32%; these peaks were then excluded for 30s within a mass window of 1.2 m/z. A tandem mass spectrum was acquired in the mass analyzer with a resolution of 60,000. The AGC target value was 1.00E+05. The ion selection threshold was 2.45E+3 counts, and the maximum allowed ion injection time was 50 ms for full scans and 120 ms for fragment ion scans. Spectra were searched against the Uniprot Mouse (Mus musculus) database using MaxQuant (2.0.3.1) with the following parameters: parent mass tolerance of 10 ppm, constant modification on cysteine alkylation, variable modification on methionine oxidation, deamidation of asparagine and glutamine. Search results were entered into Scaffold DDA software (Proteome Software, Portland, OR) for compilation, normalization, and comparison of spectral counts, etc. The filtering criteria of protein identification were with no bound false discovery rate (FDR) of protein and peptide, with 1 minimum peptide count. Protein expression levels were exported to .csv and graphs plotted using GraphPad Prism.

### GO analysis of relative protein expression

The ratio of the number of proteins expressed by a cell type in a particular GO analysis to the total number of proteins expressed by the cell types is estimated as an enrichment factor.

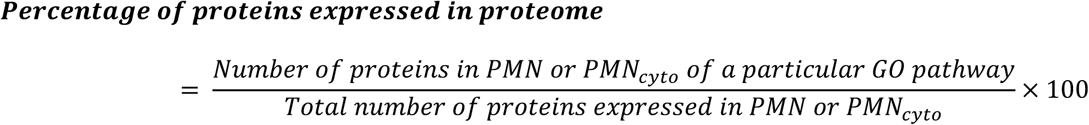

### Bacterial culture

*Pseudomonas aeruginosa* (PA-GFP-01 strain; gift from Dr. Terry Manchen, University of California, Berkeley, CA72), constitutively expressing GFP, was grown in LB Broth (Sigma L3522) with carbenicillin (Goldbio, #C-103-5). The titre of PA was calculated from colony-forming units (CFU) obtained from serial dilution of an overnight-grown culture plated on LB-carbenicillin plates.

### NETosis assay

NETosis assay, LPS-challenged/ PA-infected mice blood/ BALF were collected and centrifuged. The supernatant was used for elastase ELISA following the manufacturer’s (mouse neutrophil elastase ELISA kit; ab252356) protocol. The frequency of NET formation is plotted as the concentration of NE in blood/BALF by estimating the OD at 450 nm.

### Mouse PMN isolation

Femur and tibia bones were removed from mice immediately after euthanasia by KXA. Bones were initially kept in RPMI medium containing heparin (10 U/ml), and then flushed with HBSS-prep buffer (Ca^2+^ and Mg^2+^–free HBSS with 20 mM HEPES, 0.1% BSA). Cell suspension was filtered through a 40 mm cell strainer (Falcon 352340), centrifuged at 300g for 5 min at 40°C to pellet down. PMN were isolated by the immunomagnetic negative selection method (neutrophil isolation kit, Mitenyi Biotec, #130-097-658).

### Bronchoalveolar lavage Fluid (BALF) collection

After the termination of the experiment, mice are anesthetized. The depth of anesthesia will be checked by paw pinch to confirm the absence of pain reflex, followed by cervical dislocation. A midline incision is made along the ventral surface of the neck to expose the trachea (approximately 5mm). The trachea is cannulated with a 20-G tip needle attached to a 1 mL syringe; the lungs are instilled with sterile pyogen-free physiological saline until a total lavage volume of 2.5mL is collected.

### Bacterial killing assay

PMN and PMN_cyt_-mediated bacterial killing assay was adapted from (Gille et al., 2006; Isberg & Falkow, 1985). Constant number of PMN/ PMN_cyto_ were incubated with GFP-PA at 1:100 (cell: bacteria) ratio for 30 min for ingestion and non-phagocytosed bacteria were removed from incubation medium by gentle centrifugation at 300 rcf. Bacteria killing was assessed 30 min later. Internalized bacteria were released from PMN/ PMN_cyto_ by 1% Triton-X. The number of colonies formed was used to calculate the bacterial titer.

### Adoptive Transfer of PMN_cyto_ in mice

Donor mice are first challenged with LPS i.p 10mg/kg for 3h before being transferred 30 minutes before euthanizing the donor mice by cervical dislocation under anesthesia. Donor mice are euthanized, the lung is collected, and PMN and PMN_cyto_ are isolated and sorted. Recipient mice will be injected intratracheally with a 1×10^4^ concentration of PMN, or PMN_cyto_, or Normalized PMN_cyto_ (max 40µl). The mice are infected with PA (1×10^5^ CFU). After 24h mice were euthanized by cervical dislocation under anesthesia in BSL2 facility.

### Inflammatory lung injury evaluation in mice

Mice are anesthetized using KXA, then mice are administered EBA via retro-orbital injection 20% body weight ratio, max volume 100µl, using a 29G insulin syringe (150µl max volume). It is injected 30 minutes before sacrificing the mice by cervical dislocation, and later, the lung is harvested. The lung injury level is calculated based on the methods described previously (Wang & Lai, 2014). Followed by i.v. Evans blue (EB) [MilliporeSigma E2129, St. Louis, MO], the lung is perfused using cold PBS. Both lobes are harvested, and one lobe is weighed (wet weight) upon harvest. After measuring the wet weight, the lobe is left to dry at 60°C for 16h. Upon complete dehydration, the lung lobe is weighed again for dry weight and recorded. The other lobe is incubated at 60°C for 16h in formamide for EB extraction. After 16h of incubation, the formamide solution’s optical density (OD) is measured to estimate the quantity of EB released and recorded.

The lung injury level is measured by;

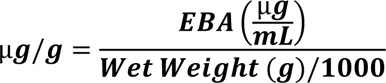

### Mitochondria isolation, transfer, and imaging

For mitochondria isolation, we followed the work published by I.R. Lanza and K.S. Nair in 2009. We prepared a homogenization buffer (KCl+Tris+MgCl_2_+ATP+EDTA) and a resuspension buffer (sucrose+KH_2_PO_4_+Mg acetate+EDTA) in the concentrations previously described (Lanza & Nair, 2009). HL60 cells were suspended in the homogenization buffer and homogenized using a Dounce homogenizer (30-35 swift movements). It is then centrifuged at 720g for 5 minutes at 4°C. The supernatant is collected (as SN#1), and the pellet is resuspended in the homogenization buffer. The homogenization step is repeated once again to increase the yield of isolated mitochondria. The supernatant is collected (as SN#2), and both SN#1 and SN#2 are centrifuged at 9000g at 4°C for 5 minutes. The pellets collected are the isolated mitochondria, and it is then resuspended in resuspension buffer for mitochondria transplantation. For mitochondrial transplantation, the PMN_cyto_ is cocultured with the isolated mitochondria for 30 minutes in an incubator at 37°C and 5% CO_2_. After 30 minutes, they are washed and centrifuged at 300g for 5 minutes at 4°C to remove free mitochondria. The stained cells (DNA-SYTO40, PMN_cyt_-Ly6G, native mitochondria-MitoTracker Green, transplanted mitochondria-MitroTracker Red CMX) were collected in a fibronectin-coated 35mm petri dish with a 14mm microwell with a No.1.5 coverglass. The cells were allowed to adhere as previously described. Images were acquired along the Z axis, using the Zeiss LSM 710 BiG 2 GaAsP detector microscope controlled by Zen software. The acquired images underwent 3D reconstruction and cropping along the XY-axis using IMARIS to maintain high resolution. Subsequently, they were converted into maximum-intensity Z-stack merges and orthogonal projections using ImageJ.

### Quantification, statistical analysis, and reproducibility of experiments

The graphs were generated using GraphPad Prism. All data were expressed as mean ± standard error. One-way ANOVA was used for pairwise comparison with multiple groups, and a Student’s t-test was used to determine statistical significance. The number of samples or mice per group, replication in independent experiments, and statistical tests are mentioned in the figure legends. For studies in mice, a significance level of 0.05 and a power of 0.9 were used to estimate the group sizes (JMP release 6 software: SAS, Cary, NC).

## Resources

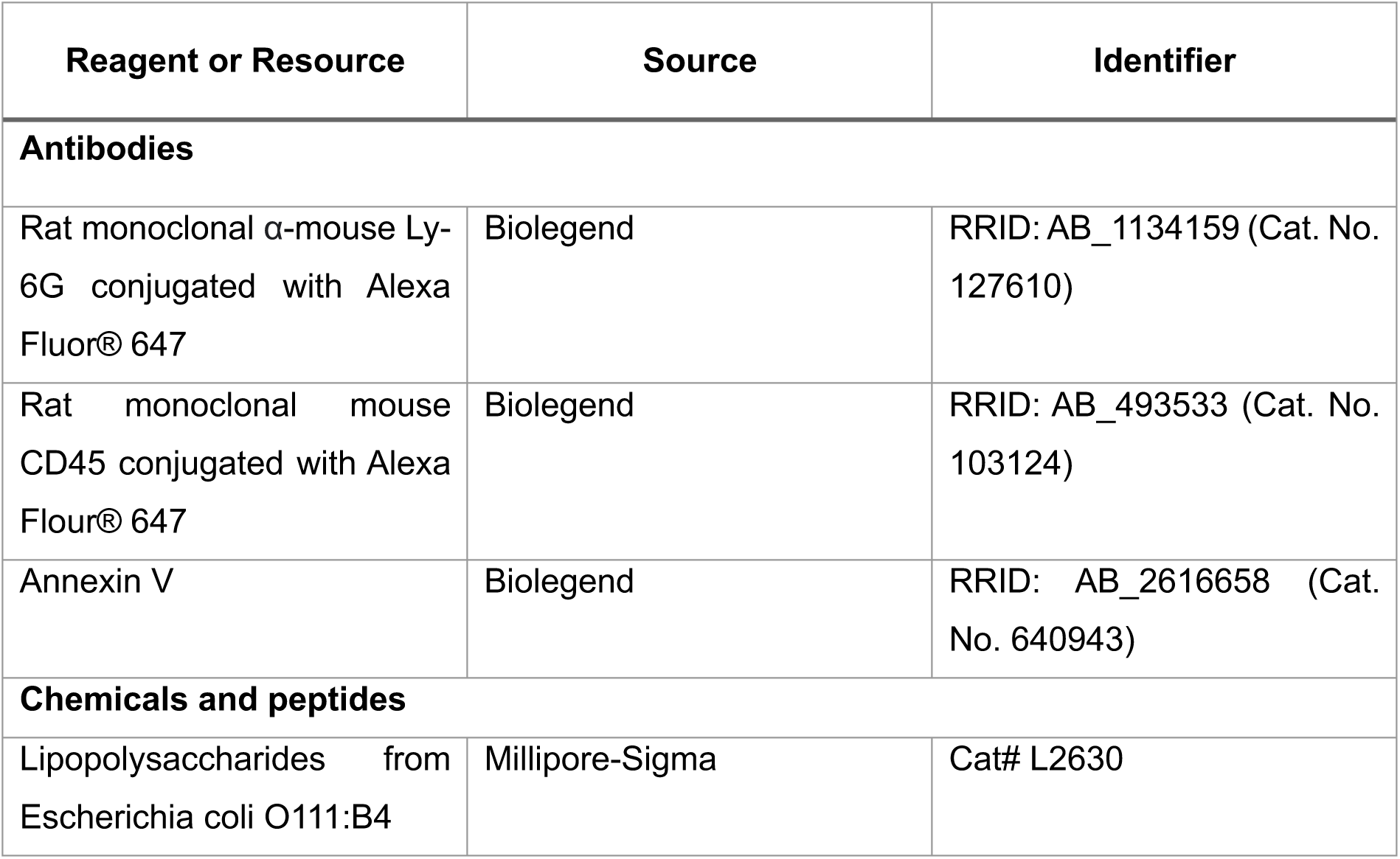
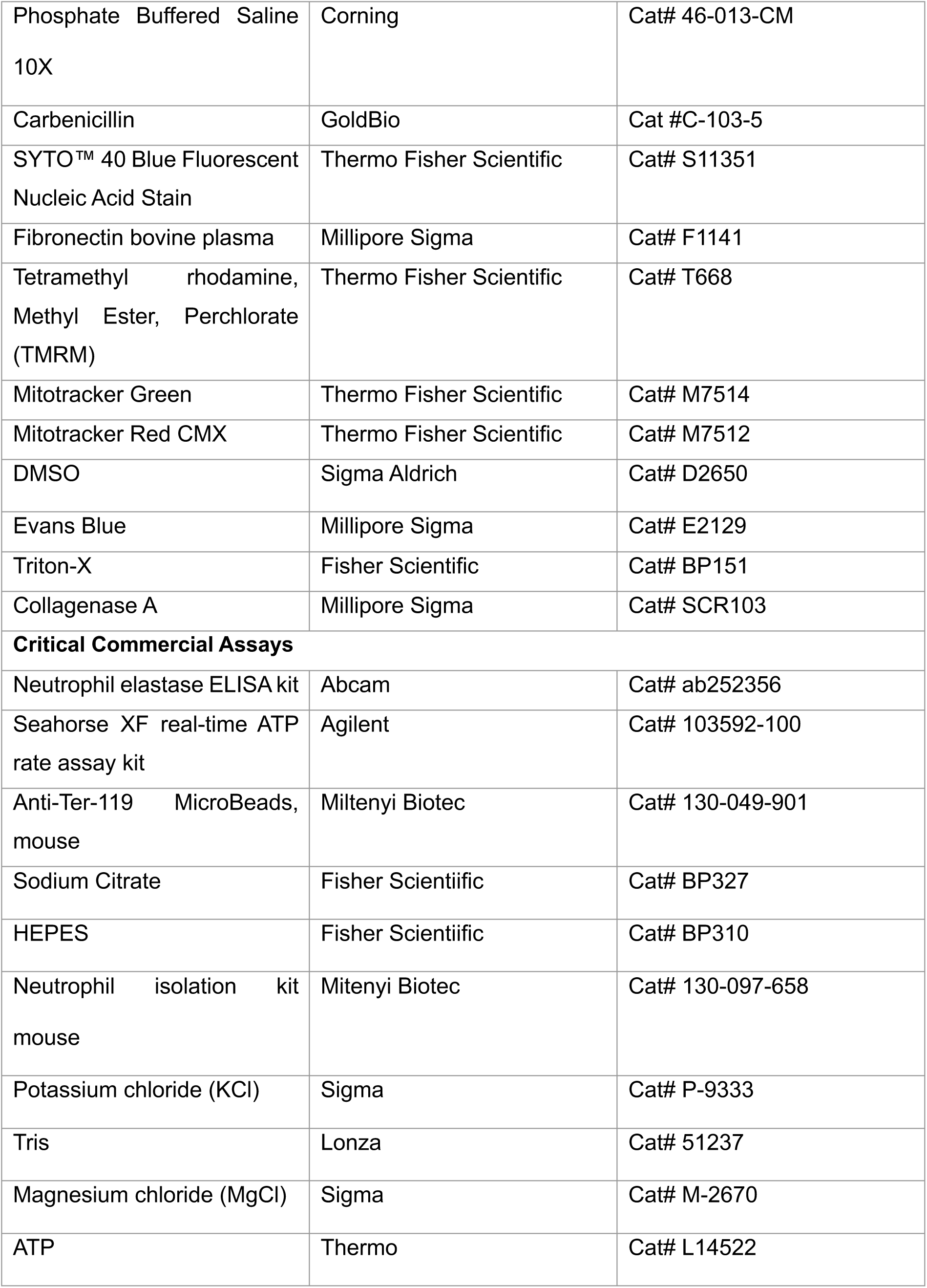
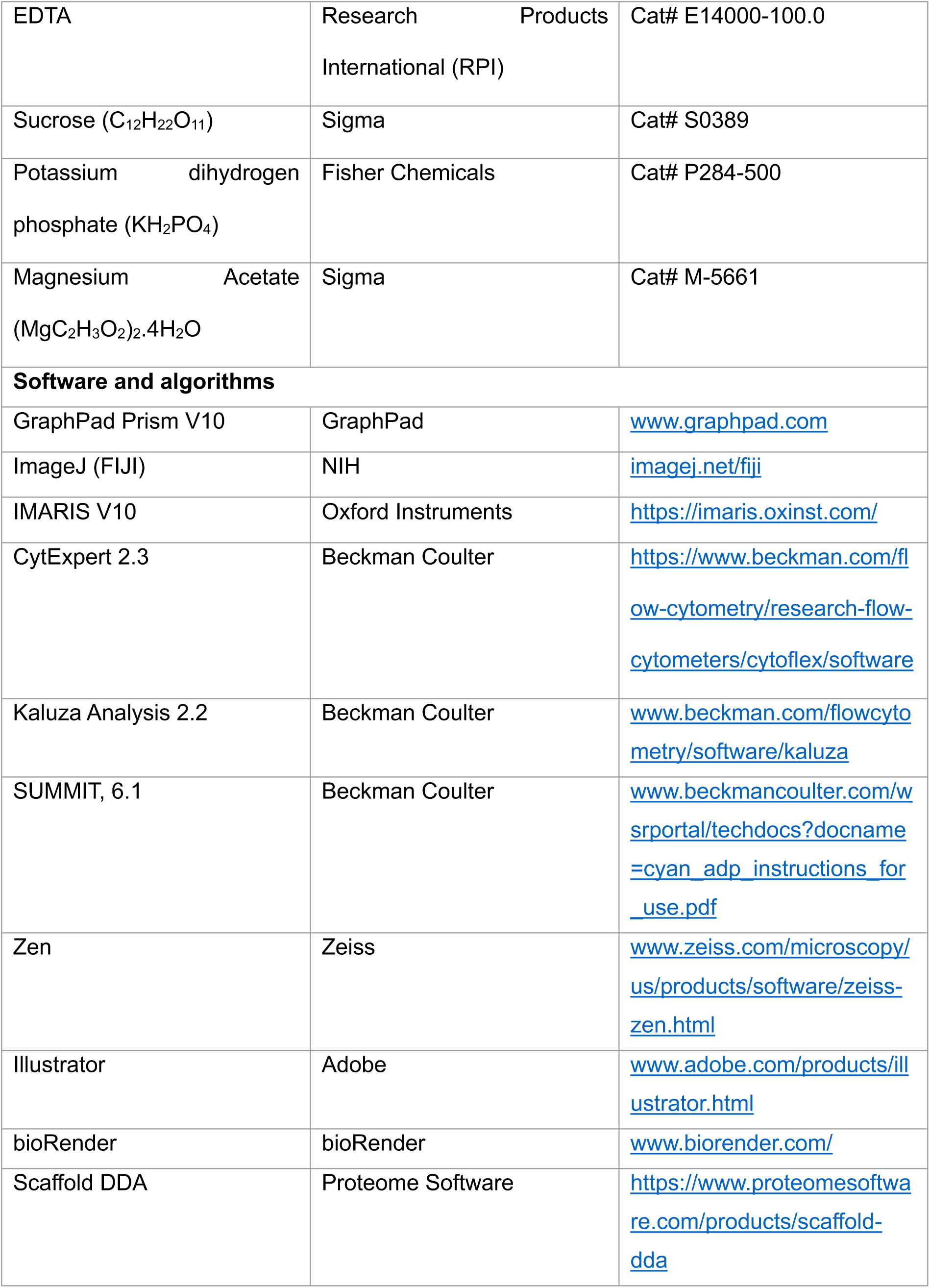
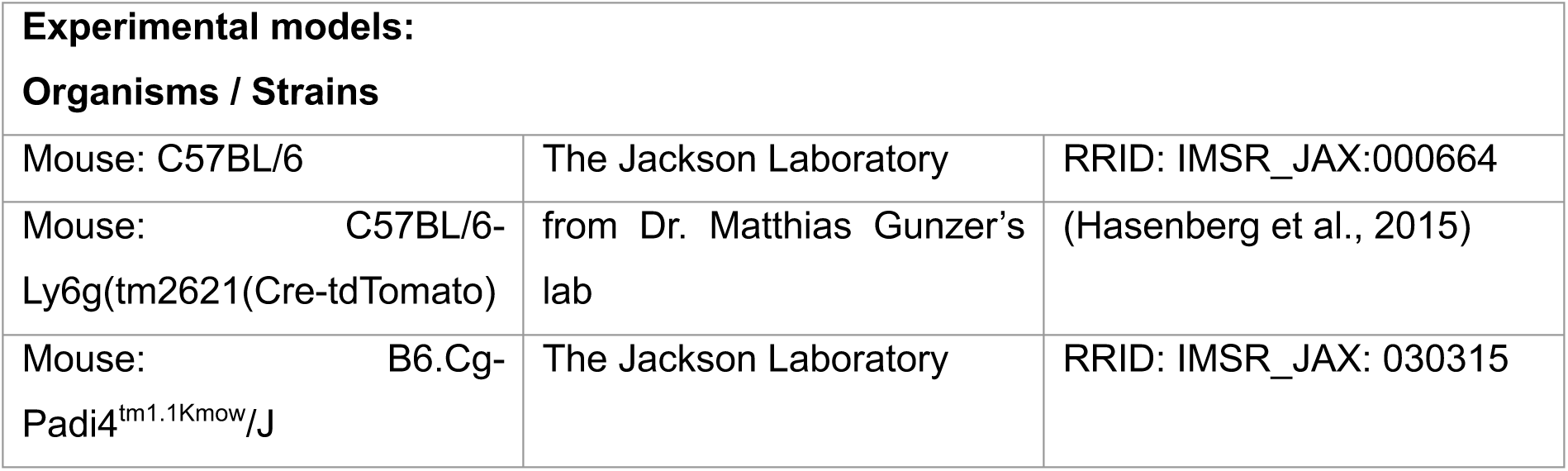

## Acknowledgement

We thank Dr. Matthias Gunzer from the University Duisburg-Essen, Essen, Germany, for providing the Catchup mice. Drs. Mohammad Anas and Dr. Joshua Thompson from the University of Illinois, Chicago, for help with bacterial cultures; the University of Illinois, Chicago Research Resource Core-Imaging facility-Dr. Peter Toth for imaging assistance; and the Mass spectrometry core facility for carrying out LC/MS analysis. This work was funded by NIH grants: 5R01HL149300, 5R01HL045638, and P01HL160469.

The work presented here represents Dr. Nithish Raj Prasad’s work carried out during his Ph.D. thesis.

## Notes

### Competing Interest Statement

The authors have declared no competing interest.

