## Supplemental Figures for "Innate Immune Function of Neutrophil Cytoplasts Generated Post-Vital NETosis"

### Supplementary Figures

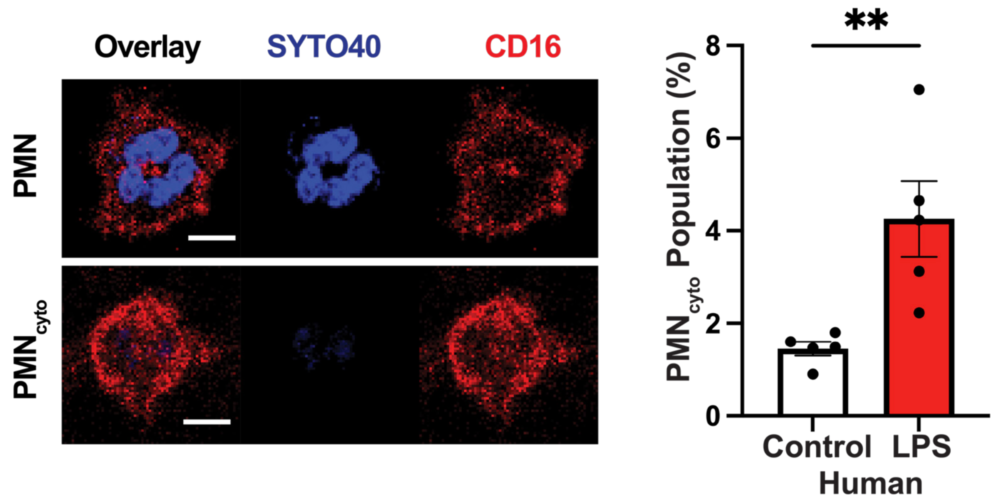


**Supplement Figure 1: Evidence for human PMN$_{cyto}$ formation.** Human pheripharal blood PMN were challenged with LPS as the same concentrations (100μg/mL) used for mouse PMN experiments (described in **Figure 1A,B**) 30m. Scale bar 5μm; N = 5; **p<0.01, shown are mean values ± SEM.

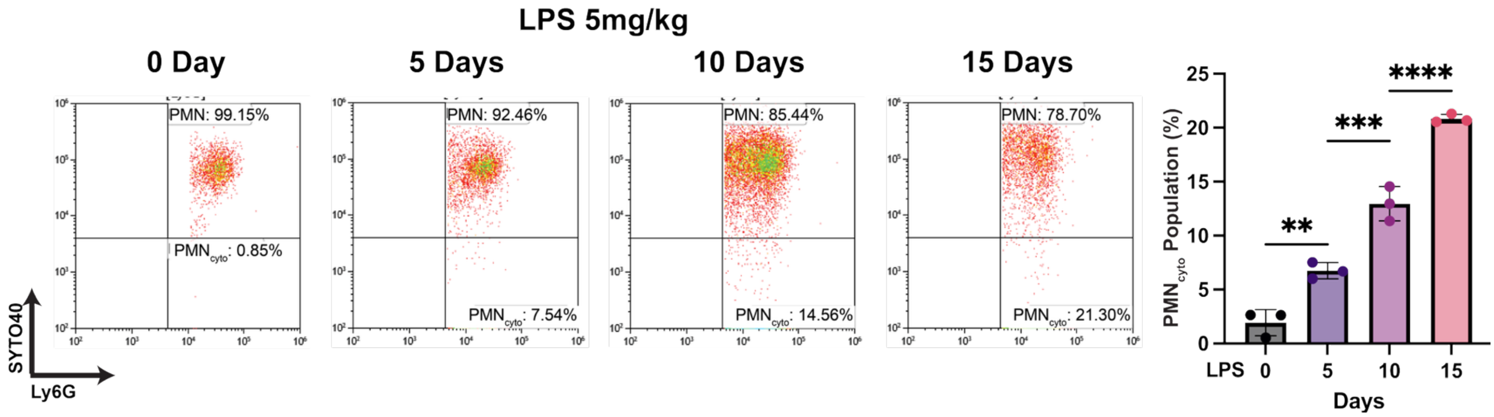


**Supplement Figure 2: Time course of PMN$_{cyto}$ generation in lungs after endotoxemia**. PMN$_{cyto}$ number was measured after challenging WT mice with LPS (5mg/kg; i.p.) for 0, 5, 10, and 15 days. Post LPS challenge, lungs were harvested, single cell suspensions were prepared, stained with SYTO40 and Ly6G-AF647, and cells were analyzed by flow cytometry. n=3; ns= not significant; *p<0.05; **p<0.01; shown are mean ± SEM.

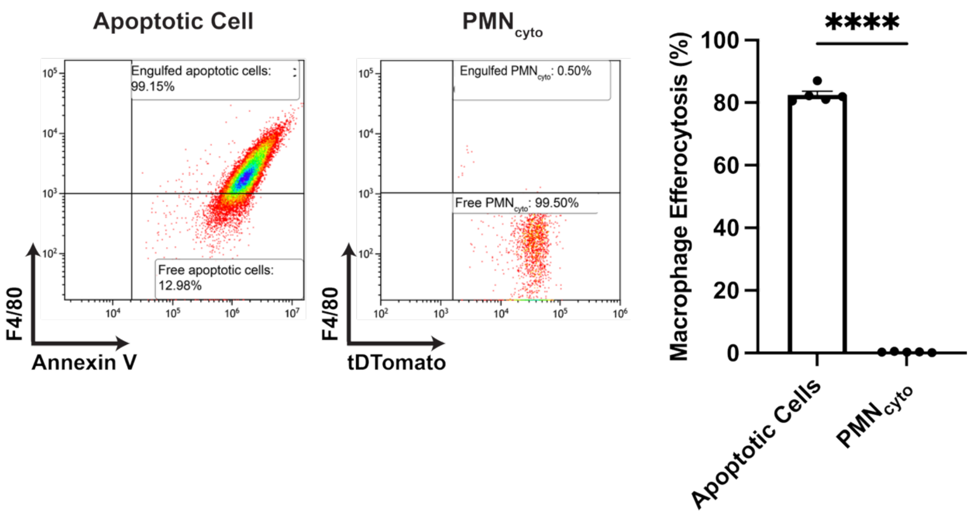


**Supplement Figure 3: Macrophages fails to clear PMN$_{cyto}$.** Mouse bone marrow macrophages incubated for 30 min with apoptotic PMN (induced by UV (Nepal et al., 2019)) stained with annexin V (red) or PMN$_{cyto}$ collected from Catchup mice 24h after LPS challenge. Quantitative analysis of macrophage efferocytosis of apoptotic PMN vs. PMN$_{cyto}$ by FACS shows ineffective clearance of PMN$_{cyto}$. Apoptotic cells stained with Annexin V; PMN$_{cyto}$ (td-Tomato) from Catchup mice; macrophages stained with F4/80. n=5, ****p<0.0001, shown are mean ± SEM.
